# Physiological and transcriptional changes associated with obligate aestivation in the cabbage stem flea beetle (*Psylliodes chrysocephala*)

**DOI:** 10.1101/2024.04.08.588545

**Authors:** Gözde Güney, Doga Cedden, Johannes Körnig, Bernd Ulber, Franziska Beran, Stefan Scholten, Michael Rostás

**Affiliations:** Agricultural Entomology, Department of Crop Sciences, University of Göttingen, Göttingen, Germany; Department of Evolutionary Developmental Genetics, Johann-Friedrich-Blumenbach Institute, GZMB, University of Göttingen, Göttingen, Germany; Population Ecology Group, Friedrich Schiller University, Jena, Germany; Division of Crop Plant Genetics, Department of Crop Sciences, University of Göttingen, Göttingen, Germany

**Keywords:** cabbage stem flea beetle, summer diapause, transcriptomics, pest biology, insect physiology, RNA-seq

## Abstract

1

Aestivation is a form of seasonal dormancy observed in various insect species, usually coinciding with the summer season. *Psylliodes chrysocephala* (Coleoptera: Chrysomelidae), the cabbage stem flea beetle, is a key pest of oilseed rape and obligatorily aestivates as adult in late summer. At present, our understanding of the physiological and transcriptional changes linked to aestivation in *P. chrysocephala* is still limited. In this study, physiological parameters and RNA-seq analyses were performed with laboratory-reared beetles at pre-aestivation, aestivation, and post-aestivation stages. Measurements of CO_2_ production supported the notion that aestivating beetles dramatically reduce their metabolic rate and, together with assessments of reproductive maturation, allowed precise discrimination between the three adult stages. Aestivating beetles showed a reduction in carbohydrate reserves and an increase in lipid reserves compared to pre-aestivating beetles, indicating that aestivation is associated with drastic changes in energy metabolism. In agreement with these findings, we found that genes involved in carbohydrate and lipid metabolism, digestion, and mitochondrial activity are differentially expressed between the three stages. Furthermore, RNA-seq analysis suggested the regulation of transcription factors associated with aestivation maintenance and the involvement of cytochrome P450s in conferring a summer-resistant phenotype during the aestivation period. In conclusion, this study represents the first exploration of the transcriptomic and physiological aspects of the aestivation response in *P. chrysocephala*.

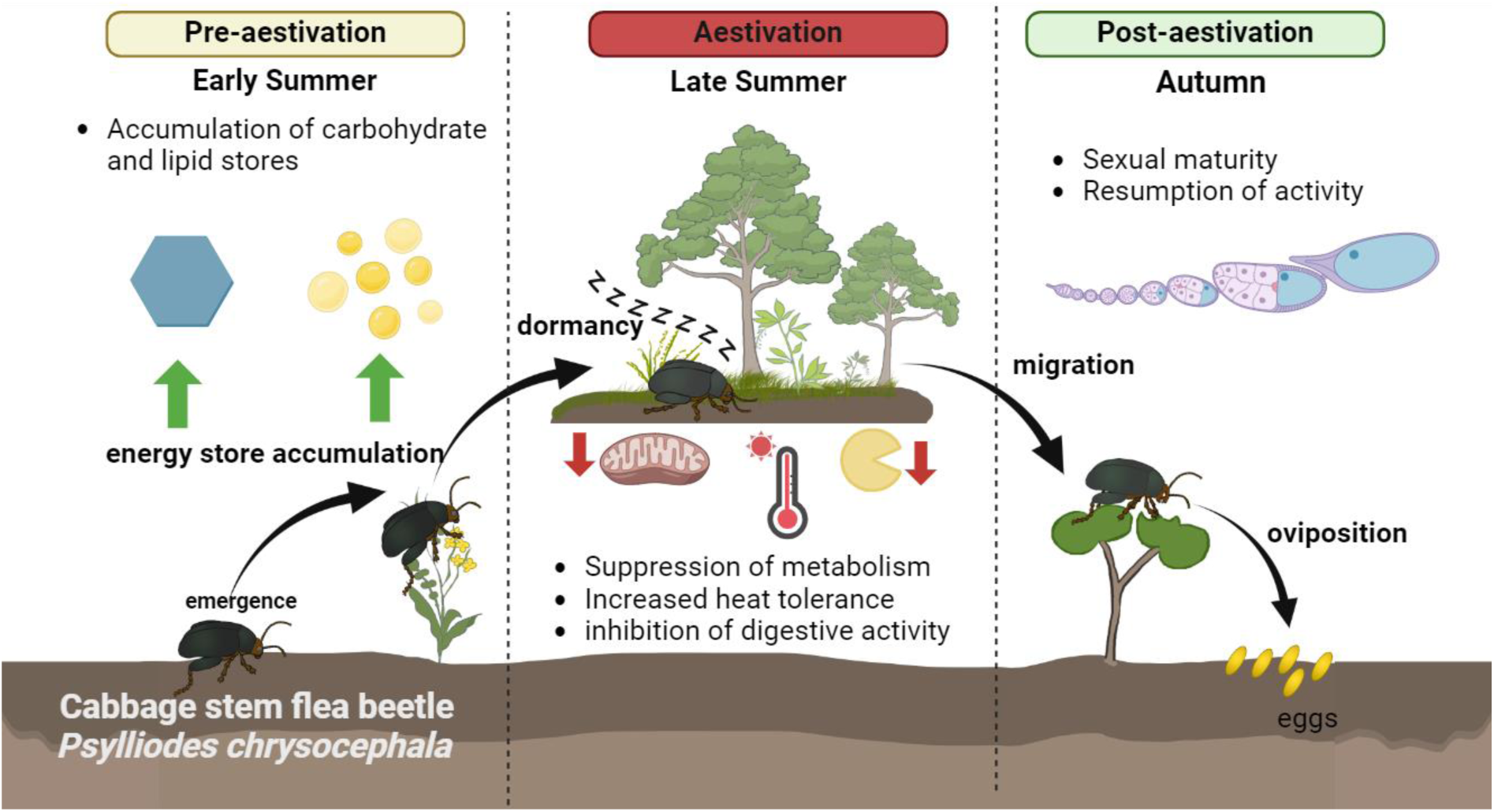

- *P. chrysocephala* obligatorily aestivate as sexually immature adults in summer
- Aestivation entails metabolic suppression, body composition changes and resistance
- Accordingly, metabolism and stress genes were differentially expressed
- The findings can support the development of innovative pest management strategies

## 2 INTRODUCTION

Diapause in insects is characterized by a predetermined cessation of development during immature stages or the absence of reproductive activity in adults. It is also characterized by a reduction in metabolic rate and behavioral activity (Denlinger, 2002; Schiesari and O’Connor, 2013). Aestivation is a type of diapause that typically occurs during the summer season as opposed to the winter hibernation observed in many temperate insect species (Saulich and Musolin, 2018; Storey and Storey, 2012). Both winter diapause and aestivation are dormant states in anticipation of unfavorable conditions and, unlike quiescence, do not occur spontaneously (Gill et al., 2017). Diapause is either induced by environmental conditions (facultative diapause) or is genetically determined (obligate diapause). Facultative diapause is prevalent among insects and is primarily triggered by token stimuli such as a decrease in photoperiod or temperature. In contrast, the less common obligatory diapause occurs at a genetically predetermined stage in the species’ life cycle, regardless of prevailing environmental conditions (Denlinger, 2022). However, the survival success of this inactive phase relies on its synchronization with unfavorable seasonal conditions, which may explain a more frequent occurrence in univoltine species (Gill et al., 2017).

Both obligate and facultative diapause are usually preceded by a diapause preparation phase in which the insect exhibits various adaptive behaviors to enhance the likelihood of survival during the subsequent dormant phase, such as migrating to protective areas and increasing feeding (Denlinger, 2002; Koštál, 2006). The feeding period allows the accumulation of energy reserves such as triglycerides and glycogen (Hahn and Denlinger, 2011; King et al., 2020), which can be catabolized to maintain energy homeostasis during diapause. For instance, *Colaphellus bowringi* adults reared under long-day conditions accumulate lipids to fuel their aestivation (Tan et al., 2017; Zhu et al., 2019). Similarly, *Leptinotarsa decemlineata* adults exposed to short-day conditions accumulate large amounts of lipids to be catabolized during winter diapause (Güney et al., 2021). Additionally, insects entering diapause may upregulate the production of protective agents, including trehalose (Elbein et al., 2003; Huang et al., 2021), glycerol (Hayward et al., 2005; Kojić et al., 2018), and heat shock proteins (Rinehart et al., 2007; Sømme, 1964) to cope with harsh environmental conditions. Furthermore, previously accumulated energy reserves can support post-diapause development, migration, or reproduction (Sinclair, 2015; Sinclair and Marshall, 2018).

A large number of RNA-seq studies have compared diapausing and non-diapausing stages of insects (Hao et al., 2019; Kang et al., 2016; Kankare et al., 2016; Lebenzon et al., 2021; Ren et al., 2018). However, studies on obligatory aestivation are scarce due to the facultative and winter-occurring nature of the majority of the diapause responses in insects. A rare example of an insect that obligatorily enters diapause during summer is *Galeruca daurica* (Coleoptera: Chrysomelidae) (Zhou et al., 2019). A transcriptomics study on *G. daurica* found evidence for the regulation of various metabolic pathways, including fatty acid biosynthesis during the transition into and out of the aestivation. (Yan et al., 2021). These results suggest that the physiological changes associated with diapause are regulated at multiple levels, albeit studies linking them are scarce.

The cabbage stem flea beetle, *Psylliodes chrysocephala* (Coleoptera: Chrysomelidae) is a major pest of winter oilseed rape (*Brassica napus subsp. napus*) in northern Europe. Adults of *P. chrysocephala* inflict damage on oilseed rape plants by feeding on leaves, while larvae mine in stems and petioles (Ortega-Ramos et al., 2022; Williams, 2010). After emerging in early summer, adult *P. chrysocephala* engages in a brief period of intensive feeding before entering aestivation until August/September, usually seeking out sheltered areas like woodlands and hedgerows (Ankersmit, 1964). Aestivating beetles neither feed nor reach reproductive maturity until aestivation is completed in 1-2 months. Subsequently, beetles invade newly sown oilseed rape fields to feed on cotyledons or young leaves and oviposit in soil cracks located near the host plant (Bonnemaison & Jourdheuil, 1954; Ortega-Ramos et al., 2022). Given the obligate nature of aestivation in *P. chrysocephala* (Sáringer, 1984), the adult stage can be categorized into three phases, namely pre-aestivation (newly emerged adults), aestivation, and post-aestivation.

To enhance our understanding of the mechanisms underlying aestivation in this economically important flea beetle species, we examined the physiological and transcriptional dynamics involved in this process. We hypothesized that aestivation is linked to various physiological and transcriptional changes that modulate cellular metabolism, alter body composition, and enhance stress tolerance under summer conditions in diapausing *P. chrysocephala*. Hence, we monitored carbon dioxide emission, energy reserves, and reproductive maturity across pre-aestivation, aestivation, and post-aestivation stages. We also identified differentially expressed genes in female beetles between these stages and conducted enrichment analyses using a functionally annotated *de novo* transcriptome assembly. Lastly, we analyzed gene expression patterns to further elucidate the findings from the enrichment analyses.

## 3 MATERIALS AND METHODS

### 3.1 Insect culture

*Psylliodes chrysocephala* was reared on potted oilseed rape plants (*Brassica napus ‘*Marathon’) in BugDorm-2120 rearing tents (60 x 60 x 60 cm; BugDorm, Megaview Science Co., Ltd., Taichung, Taiwan) as described in Cedden et al. (2024). The tents were kept under controlled conditions at 20 ± 2 °C and 65.5 ± 10% relative humidity with a 16:8 h light-dark cycle. Light was provided by daylight spectrum LED lamps (Bioledex GoLeaf E2, DEL-KO). To obtain insects at different ages, newly emerged adult beetles were kept in groups of 15-20 individuals in ventilated PET boxes (11 x 6 x 15 cm) with detached oilseed rape leaves under the conditions described above.

### 3.2 Carbon dioxide emission

Carbon dioxide emission rates (VCO_2_) of female and male beetles (n = 16 per sex) were recorded using an LI-820 VCO_2_ Gas Analyzer (LI-COR Environmental GmbH, Bad Homburg, Germany) at 5-day intervals for 60 days following adult emergence. Details of the measurement process are described in Cedden et al. (2024). The weight corrected ΔCO_2_ ppm values were analyzed using two-way ANOVA followed by Bonferroni’s multiple comparison test (GraphPad Prism v10.0).

### 3.3 Reproductive maturation

Ovaries were dissected from 5, 25, and 45 days old females (n = 8) in phosphate-buffered saline (7.4 pH) to assess morphological changes. Dissected ovaries were examined under a Leica TL3000 Ergo stereo microscope (Microsystems, Switzerland) and photographed using a Leica DMC5400 camera. The area of each ovariole per female was measured using ImageJ (v1.53t, https://imagej.nih.gov/ij/index.html). The average ovariole areas per female were statistically analyzed using one-way ANOVA followed by Tukey’s test (GraphPad Prism v10.0).

Oviposition activity was investigated by placing newly emerged individual female and male beetles (n = 14 pairs) into plastic boxes containing fresh oilseed rape leaf discs that were replenished every two days. The adult pairs were monitored daily for the presence of eggs to reveal the beginning of the oviposition period. Data analysis involved plotting a Kaplan-Meier curve, with the laying of the first eggs serving as the endpoint. The median female age at first oviposition was calculated using GraphPad Prism v10.0.

### 3.4 Survival under heat stress

Pre-aestivating adults (initially 5 days old) and aestivating adults (initially 30 days old) were kept at 30 °C and 60% relative humidity, maintaining the regular 16:8 h light-dark cycle. Survival was monitored daily for 6 days (n = 30). Fresh leaf discs (approx.130 mm^2^) punctured from the first true leaves of randomly selected oilseed rape plants (growth stage BBCH 30-35) were provided to both groups every two days. Survival curves were plotted using the Kaplan–Meier method. The *P* value and hazard ratio (hazard in pre-aestivation vs. hazard in aestivation) were calculated by the log-rank test using GraphPad Prism v10.0.

### 3.5 Composition of energy reserves and water content

Body composition analyses were performed to determine the contents of energy reserves in 5, 30, and 55 days old female and male beetles (n = 6-10 per sex). At the designated time points, the beetles were frozen with liquid nitrogen and kept at −80°C following fresh weight measurements. Total protein, total lipid, glycogen, and soluble carbohydrate contents were measured using a previously established method by Foray et al. (2012) with minor modifications described in Sporer et al. (2021). This method enables the quantification of major energy reserves in individual beetles. In addition, water contents were assessed using another batch of female and male beetles (n = 10 per sex) by first measuring individual fresh weights before freezing them with liquid nitrogen and letting the samples dry at 60 °C for 72 h to measure the dry weights, which were subtracted from the initial fresh weights. Measurement results for each content were analyzed using two-way ANOVA, with one of the factors being sex and the other being the adult stage (Table S1).

### 3.6 Library preparation

Total RNA from pre-aestivating (5 days old), aestivating (30 days old), and post-aestivating (55 days old) female beetles were extracted using the Quick-RNA Tissue/Insect Kit (Zymo Research, Freiburg, Germany). Subsequently, the RNA samples were purified using the TURBO DNA-free™ kit (Thermo Fisher Scientific, Waltham, MA, USA). To eliminate sex-related variability, only female beetles were sampled. RNA integrity was verified using an Agilent 2100 Bioanalyzer and an RNA 6000 Nano Kit (Agilent Technologies, Santa Clara, CA, USA). RIN values ≥ 7.0 were considered appropriate for mRNA library preparation. In total, 10 libraries (4, 3, and 3 libraries respectively per pre-aestivation, aestivation, and post-aestivation stages) were prepared using NEBNext® Poly(A) mRNA Magnetic Isolation Module followed by NEBNext kit (E7490, NEB, Ipswich, MA, USA). The quality and quantity of the libraries were checked on an Agilent 2100 Bioanalyzer using the Agilent DNF-935 Reagent Kit (Agilent Technologies, Santa Clara, CA, USA). Libraries were pooled to equimolar amounts, and a total concentration of 3.4 ng/µL was obtained. Sequencing was performed by BGI Genomics Tech Solutions Co. Ltd (Hong Kong) on a DNBSEQ-T7 platform. Raw read files were deposited in the Sequence Read Archive (SRA) database of NCBI under the accession numbers: SAMN33022552 - SAMN33022561.

### 3.7 *De novo* assembly and functional annotation

Erroneous k-mers from paired-end reads were removed using Rcorrector (v1.0.5) with default options (Song and Florea, 2015), and unfixable reads were discarded using the “FilterUncorrectabledPEfastq.py” function in Transcriptome Assembly Tools (Song and Florea, 2015). Adaptor sequences were removed, and reads with a quality score greater than 30 were retained using TrimGalore! (v0.6.7). The cleaned reads (n = 3 per three adult phases) were assembled *de novo* using Trinity with default options. In total, 224 million bases were assembled covering 341,670 transcripts, including putative isoforms (https://doi.org/10.6084/m9.figshare.21922938). The d*e novo* assembly had an N50 value of 1532 and a BUSCO (v5.4.2) completeness score of 96.7% when compared against the Endopterygota lineage (BUSCO.v4 datasets). Putative isoforms were combined to obtain a supertranscriptome containing 189,229 transcripts in total. The supertranscriptome was deposited at GenBank as a Transcriptome Shotgun Assembly (TSA) under the accession number GKIH00000000.1.

The transcriptome (including isoforms) was annotated using Trinotate (v3.2.2). This tool combines the outputs of various packages, such as NCBI BLAST+ (v2.13.0; nucleotide and predicted protein BLAST), TransDecoder (v5.5.0; coding region prediction), SignalP (v4.0; signal peptide prediction) (Petersen et al., 2011), TmHMM (v2.0; transmembrane domain prediction) (Krogh et al., 2001), and HMMER (v3.3.2; homology search) into an SQLite annotation database. The uniprot_sprot (04/2022) and pfam-A (11/2015) databases were downloaded using Trinotate and the default E-value thresholds were used during the searches with BLAST+ and HMMER. The obtained annotation database was used to extract gene ontology (GO) terms associated with individual genes using the “extract_GO_assignments_from_Trinotate_xls.pl” whereas the signals and TmHMM outputs were manually extracted using Excel spreadsheets. The longest protein-coding regions in the supertranscriptome data predicted by TransDecoder were subjected to Kyoto Encyclopedia of Genes and Genomes (KEGG) pathway annotation via GhostKoala v2.2 (https://www.kegg.jp/ghostkoala/). The annotation database was made publicly available on Figshare (https://doi.org/10.6084/m9.figshare.21922938).

### 3.8 Differentially expressed genes

Read counts per transcript were calculated using Salmon (v1.9) by mapping the cleaned reads onto our *de novo* transcriptome. Transcripts with less than 15 total read counts across all 10 libraries were discarded from further analysis. The R package “DeSeq2“(v4.2) was used to identify the differentially expressed transcripts in the aestivation vs. pre-aestivation and aestivation vs. post-aestivation comparisons. Transcripts with *P* values below 0.05 and Log_2_ Fold change (LFC) values below −1.0 or above 1.0 were accepted as significantly down- and upregulated.

### 3.9 Enrichment analyses

The “enricher” function in the R package “ProfileClusterer” was used to analyze the enrichment status of GO terms and KEGG pathways associated with the differentially expressed transcripts in the two pair-wise comparisons. All the transcripts with at least one GO annotation served as the background (12,410 transcripts). We did not distinguish between up- and down-regulates genes during the enrichment analyses due to the ambiguous nature of the GO term annotations. We plotted the top 35 most significantly enriched GO terms. (full enrichment results are available at https://doi.org/10.6084/m9.figshare.24085815).

The dataset was also investigated for the number of genes predicted with signal peptides, transmembrane domains, both, or neither. The number of genes belonging to each category was determined by manually investigating the SQLite annotation database, and Chi-squared tests were performed to compare the proportion of each category among differentially expressed genes with that among the background genes. Here, the upregulated and downregulated genes were separately analyzed, and Bonferroni correction was applied (*P* < 0.05/18 = 0.002).

Gene hits from significantly enriched GO terms coinciding with our physiological measurements were selected for the visualization of their expression pattern across the three adult stages. R v4.3 was used to Z-normalize the expression values and GraphPad Prism v10.0 was used to construct the heat maps. The names of the genes were extracted from the annotation database constructed in this study and the names were curated where necessary.

## 4 RESULTS

### 4.1 Physiological characterization of aestivation

#### 4.1.1 Metabolic rates

To estimate the timing of aestivation in our laboratory *P. chrysocephala* population, we measured the metabolic rate (ppm mg^-1^ min^-1^) of female and male adults every 5 days from day 5 to day 60 post-emergence. The metabolic rate changed significantly over time, with the day factor alone explaining 79.5% of the variation (Fig. 1A). In addition, the metabolic rate of males was significantly higher than that of females (*P* < 0.001), but there was no significant interaction between the sex and day factors (*P* = 0.967). Metabolic rate decreased between days 5 and 15, remained stable until day 40, and increased thereafter. These results confirm that *P. chrysocephala* adults undergo an obligatory aestivation phase that lasts for about 25 days.

**Figure 1.**
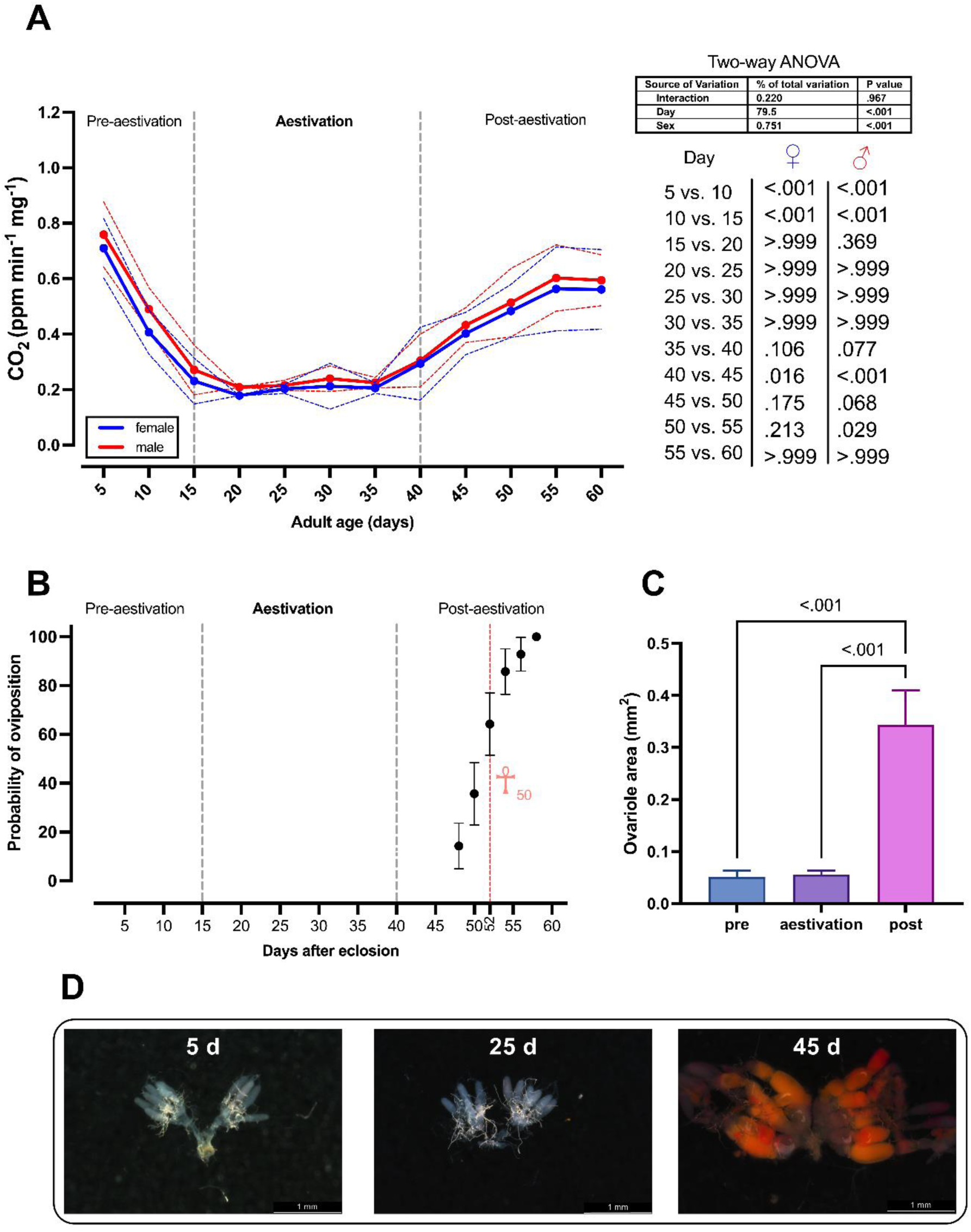
Characterizing the aestivation period in *P. chrysocephala*. (A) VCO_2_ production (ppm min^-1^ mg^-1^) of *P. chrysocephala* females (blue) and males (red) over 60 days at 5-day intervals following adult emergence. The line graph depicts the mean VCO_2_ production for each sex, with dotted lines indicating the standard deviation (n = 16 per sex). A two-way ANOVA with the factors of day and sex was performed. The table presents the percentage of explained variation and *P* values for day, sex, and their interaction. Bonferroni’s multiple comparison tests were employed to compare VCO_2_ production at each time point with the subsequent time point. Adjusted *P* values are given for females and males separately. (B) Cumulative probability of oviposition in female beetles, when paired with male beetle’s post-adult emergence (n = 14, one pair per Petri dish). Kaplan-Meier analysis determined the day at which half the population is expected to have started oviposition (☥_50_). The error bars indicate the standard error of the probability (C) Mean ± SD ovariole areas (mm^2^) in pre-aestivating (pre; 5 days old), aestivating (25 days old), and post-aestivating (post; 45 days old) females (n = 8). Tukey’s test compared ovariole areas, with significant differences (*P* ≤ 0.05) indicated above bars. (D) Representative images of ovaries examined at different time points following adult emergence (n = 8).

#### 4.1.2 Reproductive maturation

Subsequently, we investigated the reproductive maturity of females by analyzing the onset of oviposition under our laboratory conditions and by comparing the morphology of the reproductive system of females in the pre-aestivation, aestivation, and post-aestivation phases. The onset of oviposition ranged from day 48 to day 60 after emergence in the post-aestivation phase. The median onset of oviposition, i.e. the time when half of the female population had started oviposition, was estimated to be day 52 (Fig. 1B). In agreement with this observation, we found that the ovariole area was significantly larger in post-aestivating females compared to pre-aestivating and aestivating females (*P* < 0.001), while no clear morphological differences were found between pre-aestivating and aestivating females (Fig. 1C and D). In contrast to females, no clear morphological changes were observed in the reproductive system of males between the pre-aestivation, aestivation, and post-aestivation phases. (Fig. S1).

#### 4.1.3 Survival under heat stress

To test whether pre-aestivating and aestivating adults are differentially affected by high temperature, we exposed pre-aestivating and aestivating adults to 30 °C and 60% relative humidity. As expected, aestivating adults survived significantly better than pre-aestivating adults (*P* = 0.009; Fig. 2). The hazard ratio between pre-aestivating and aestivating was 2.7, indicating that pre-aestivating adults are more strongly affected by high temperature than aestivating adults.

**Figure 2.**
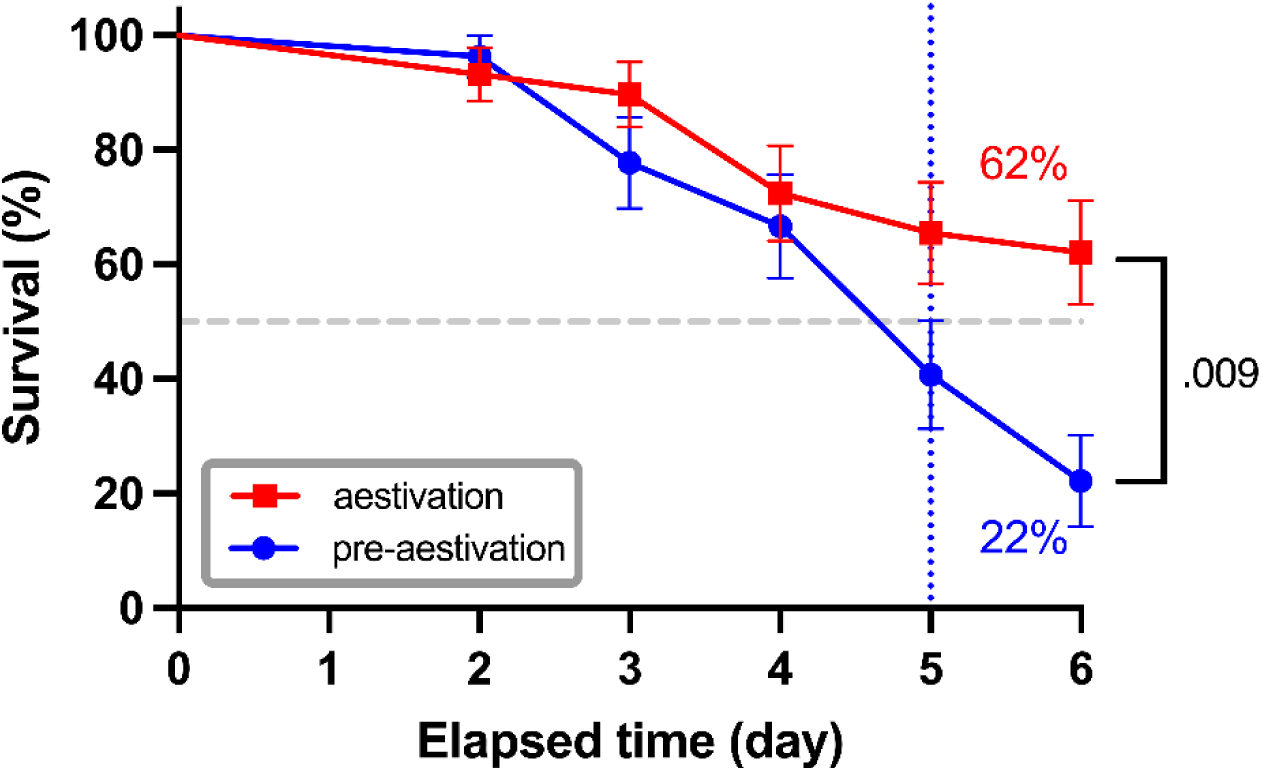
Survival of pre-aestivating and aestivating *Psylliodes chrysocephala* adults at 30°C. Pre-aestivating and aestivating adults were reared at 30°C and 60% relative humidity to simulate heat stress. Survival was monitored daily. The survival curves show the mean percentage of surviving adults ± standard error (n = 30) calculated by the Kaplan–Meier estimator. The *P* value was calculated by log-rank test. The blue vertical line indicates the median lethal time of the pre-aestivating adults.

#### 4.1.4 Composition of energy reserves and water content

To further characterize the physiological changes associated with aestivation in *P. chrysocephala*, we compared the allocation of energy reserves and water content of pre-aestivating, aestivating, and post-aestivating males and females (Fig. 3A). Protein concentration did not differ between the three stages in females but was significantly lower in post-aestivating males than in pre-aestivating males (*P* < 0.04).

**Figure 3.**
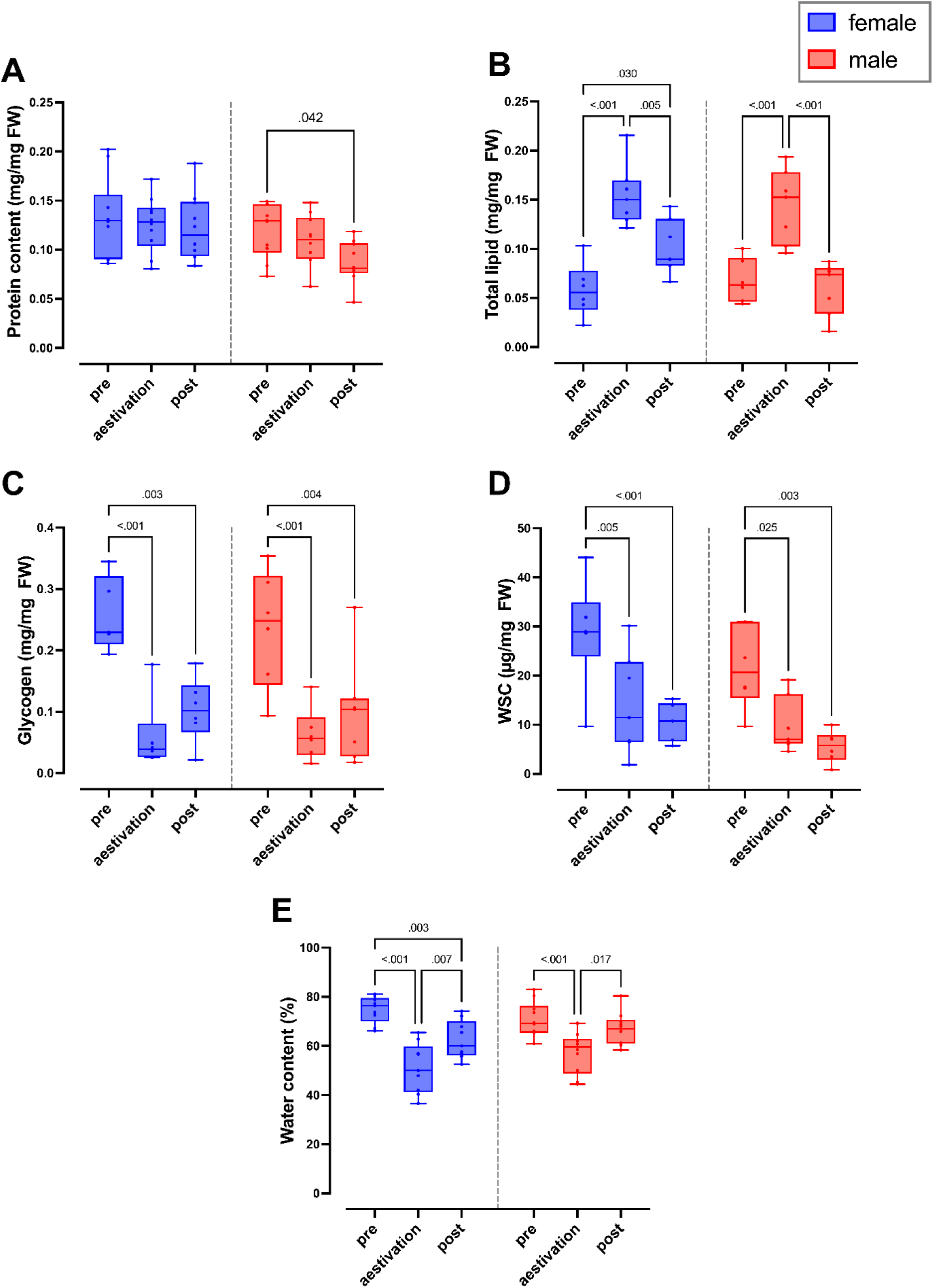
Total protein (A), total lipid (B), glycogen (C), and water-soluble carbohydrate WSC (D), and water content (E) contents of pre-aestivating (pre; 5 days old), aestivating (30 days old), and post-aestivating (post; 55 days old) *P. chrysocephala* females (blue) and males (red). Each content was divided by the fresh weight (FW) of the associated beetle and plotted as a box plot which shows the median (middle line), upper and lower quartiles (the box), and the minimum and maximum (whiskers) of the consumption values (n = 6-10 per sex). Two-way ANOVA was conducted to test whether the factors stage (pre, aestivation, and post) and sex significantly affected each measured parameter (see Tab. S1). The effect of the factor stage within sex was further investigated by Tukey’s test (*P* ≤ 0.05).

In contrast to protein levels, lipids, glycogen, and water-soluble carbohydrate (WSC) concentrations differed significantly between aestivating and non-aestivating phases in both sexes. While lipid concentrations were significantly higher during aestivation compared to the pre-aestivation phase (*P* < 0.05), glycogen and WSC concentrations were significantly lower. In the post-estivation phase, the lipid concentrations were significantly lower, whereas the glycogen and WSC concentrations remained at a similar level as in the pre-estivation phase (Fig. 3B, 3C, 3D). In addition, we observed a significantly lower WSC concentration in males compared to females, whereas protein, lipid, and glycogen concentrations did not differ between both sexes.

Water content was significantly lower in aestivating beetles compared to pre- and post-aestivating beetles. In addition, post-aestivating females had a significantly lower water content than pre-aestivating females (Fig. 3E). Taken together, our results show that aestivation is associated with drastic changes in the composition of energy reserves and water content in *P. chrysocephala*.

### 4.2 Transcriptional characterization of aestivation

#### 4.2.1 Differentially expressed transcripts

Using our RNA-seq reads and *de novo* transcriptome assembly, we performed two pairwise comparisons; aestivation vs. pre-aestivation and aestivation vs. post-aestivation. (Fig. S2). The latter stage (the stage after “vs.”) was taken as the reference for each comparison. The aestivation vs. pre-aestivation comparison revealed that 3013 and 3223 transcripts were up- and down-regulated, respectively, among the 34,368 transcripts that passed the pre-filtering. Similarly, 2,762 and 3,876 transcripts were found to be up- and downregulated, respectively, in the aestivation vs. post-aestivation comparison. Hence, about 5% of transcripts were differentially expressed depending on the adult stage. To determine the transcripts that were specifically up- or down-regulated during aestivation, we checked the overlap between the two comparisons. The overlap analysis resulted in 246 and 657 transcripts that were significantly upregulated and downregulated in aestivation compared to pre- and post-aestivation stages, respectively, indicating a bias toward down-regulation (2.7-fold compared to up-regulation) in this filtered dataset.

#### 4.2.2 Enrichment analyses

The top enriched GO terms in the aestivation vs. pre-aestivation comparison (emphasizing the initiation of aestivation) were mainly related to cellular metabolism (Fig. 4A). The analysis strongly suggested alterations in carbohydrate metabolism and mobilization. Additionally, various terms suggested alterations in mitochondrial activity, e.g., mitochondrial electron transport, tricarboxylic acid cycle, and mitochondrial respiratory chain complex I and IV terms. The most significantly enriched GO term was digestion, and various other terms and pathways related to this process, such as cellulose catabolic process and polygalacturonase, were also enriched. Another significantly enriched term related to metabolism was the fatty acid biosynthetic process. As expected, juvenile hormone (JH) esterase activity was also significantly enriched. Interestingly, terms related to chitin binding and structural constituents of cuticle were among the top enriched GO terms, suggesting potential modifications in integument or peritrophic matrix composition. Changes in molecular functions involving iron atoms were also supported by two enriched terms.

**Figure 4.**
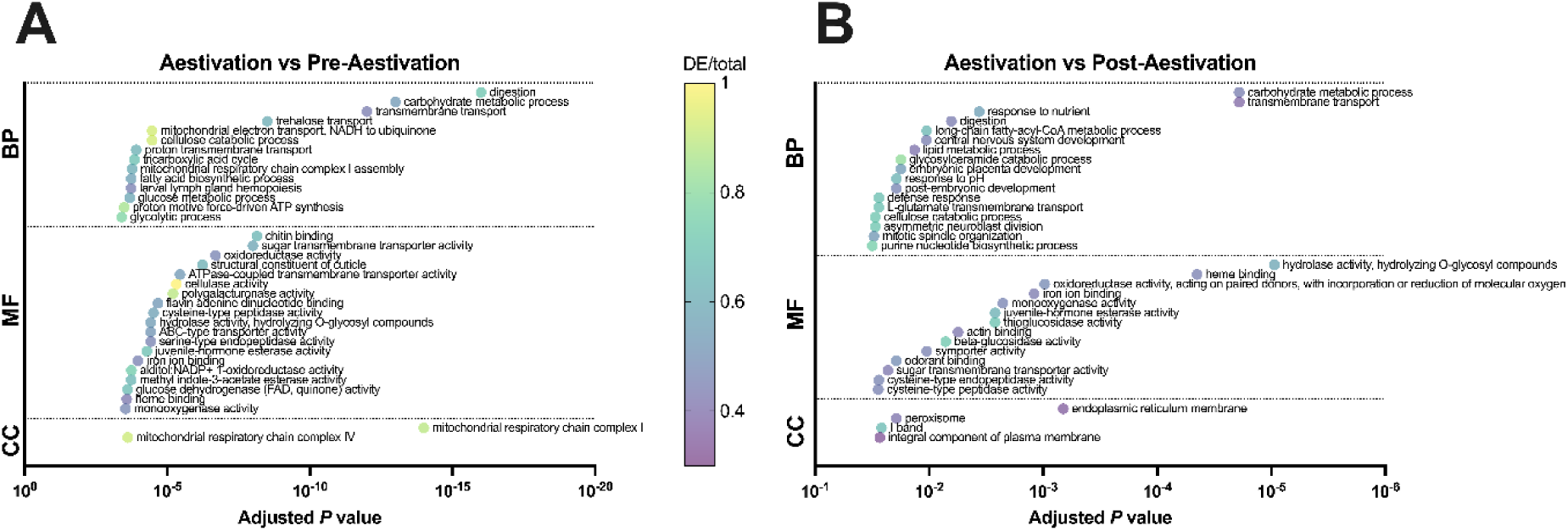
GO term enrichment results. The bubble plots of the enrichment analysis of genes identified as significantly up- or downregulated in the aestivation vs. pre-aestivation (A) and aestivation vs. post-aestivation (B) comparisons. The top 35 significantly enriched GO terms were shown and the terms were grouped into biological process (BP), molecular function (MF), and cellular component (CC). The X-axis indicates the adjusted *P* values of the enriched terms while the color of the bubble indicates the ratio between the differentially expressed (DE) genes and the total genes associated with each GO term.

The comparison between aestivation vs. post-aestivation revealed that regulation of digestive processes, cellular metabolism, JH esterase, and iron binding activity might be associated with the termination of aestivation as well (Fig. 4B). Nonetheless, distinct terms were also enriched in this comparison. Embryonic placenta development and post-embryonic development terms are likely to be related to the reproductively mature state of post-aestivating females. Interestingly, response to pH and defense response were among the top enriched biological processes. Compared to the pre-aestivation vs. aestivation comparison, we observed more terms of cellular component ontology among the top 35 enriched GO terms. These included components such as endoplasmic reticulum membrane and peroxisome, implicating their involvement in lipid metabolism and stress response. Additionally, the emergence of the term I band, which is a muscle-related structure, hints at structural changes in muscle tissue during aestivation termination.

Using the data from pair-wise transcriptome comparisons, we investigated whether there is any bias towards genes with predicted signal peptides and/or transmembrane helices among differentially expressed genes compared to the background. The analysis revealed a significant enrichment of genes containing signal peptides, transmembrane helices, or both among genes that were downregulated in both the aestivation vs. pre-aestivation and aestivation vs. post-aestivation comparisons (Fig. S3). All three categories were also significantly enriched among the genes upregulated in the aestivation vs. post-aestivation comparison.

#### 4.2.3 Investigations at the gene level

To enhance clarity in interpreting enrichment analysis outcomes, such as discerning whether biological processes are activated or deactivated, we employed heat maps. The heat maps show the expression patterns of the top 20 differentially expressed transcripts that were associated with the enriched GO terms of interest (i.e., significantly enriched and related to the physiological measurements). We also included transcripts specifically regulated during aestivation to gain further insights (Fig. 5). The selected terms were associated with carbohydrate and lipid metabolic processes, mitochondrial respiratory chain, reproduction (embryonic development and female reproductive system terms), heme binding and feeding activity (digestion and response to nutrients terms), as determined by our annotation database.

**Figure 5.**
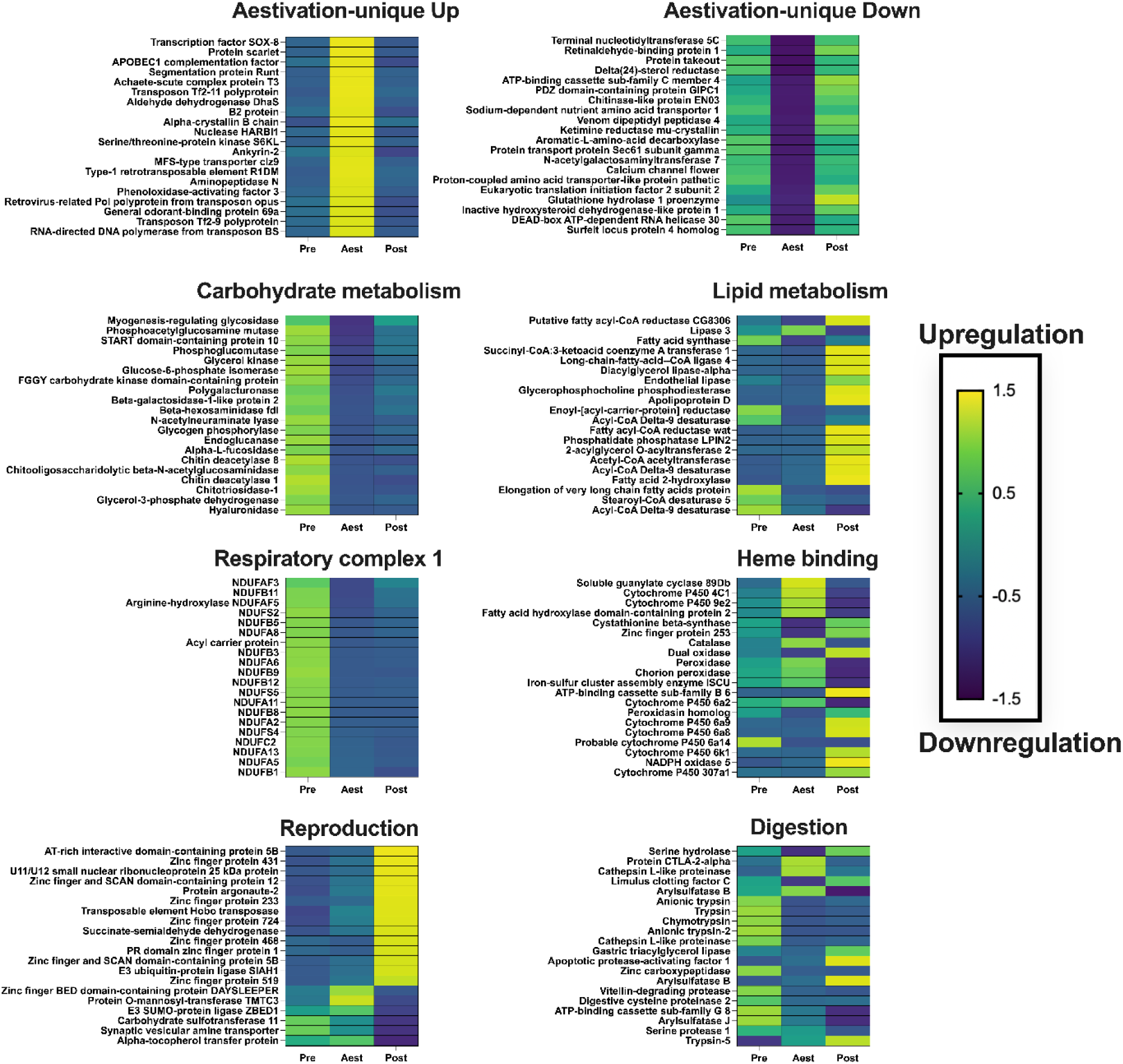
Heat maps illustrating the expression of top differentially expressed transcripts during pre-aestivation (Pre), aestivation (Aest), or post-aestivation (Post) stages of female *Psylliodes chrysocephala*. Each heatmap shows the top 20 differentially expressed transcripts from sets uniquely up- or downregulated during aestivation or the GO terms of interest from Fig. 4. Each row represents a transcript with its annotation taken from our annotation database and curated where necessary (see https://doi.org/10.6084/m9.figshare.21922938 for the original automated annotations). The color of the cells represents the z-score normalized read count values averaged per stage; dark blue indicates down-regulation and yellow cells indicate up-regulation.

Prominent among the transcripts uniquely upregulated during aestivation was a transcription factor identified as Sox-8, which might be required for the gene expression pattern associated with aestivation (Fig. 5). Additionally, a putative pigmentation protein (protein scarlet) and an mRNA editing transcript were among the top transcripts specifically regulated during aestivation. The former may contribute to increased protection against summer conditions, while the latter might facilitate intricate regulation to finely tune the aestivation transcriptome. We also noticed multiple transposon-related transcripts to be specifically upregulated during aestivation, which might be due to the compromised transposon control mechanisms linked to aestivation. The top transcripts distinctly downregulated during aestivation were associated with a range of putative functions, including protein transport and translation initiation.

The top transcripts associated with carbohydrates were especially upregulated during pre-aestivation and then downregulated during both aestivation and post-aestivation (Fig. 5). This finding is consistent with our body composition measurements. Both glycogen and WSC contents were abundant only during the pre-aestivation stage and largely absent during the following two stages.

On the other hand, important transcripts related to lipid metabolism showed a completely different pattern compared to those linked with carbohydrate metabolism (Fig. 5). Many of these transcripts, such as apolipoprotein (lipid carriers in hemolymph) and diacylglycerol lipase were specifically upregulated during the post-aestivation stage. In contrast, the transcript related to lipogenesis, fatty acid synthase, was upregulated only during pre-aestivation, presumably to accumulate the lipid reserves that peaked during aestivation according to our total lipid measurements. Also, an aestivation-specific lipase was identified, possibly responsible for energy and metabolic water production during aestivation through the catabolism of lipids.

As expected from the CO_2_ measurements, transcripts associated with respiratory chain complex I were upregulated during pre-aestivation and dramatically downregulated during aestivation, indicating metabolic suppression in the latter (Fig. 5). Interestingly, expression levels did not return to pre-aestivation levels during post-aestivation, suggesting that post-aestivating adults are not as metabolically active as pre-aestivating adults, consistent with our VCO_2_ measurements.

Among the top differentially expressed transcripts with heme binding annotation, many cytochrome P450s were identified, including 4C1. Additionally, the expressions of two peroxidases peaked during aestivation, indicating the activation of the antioxidant defense system to support the maintenance of aestivation.

As expected, differentially expressed transcripts related to reproduction were upregulated in post-aestivation (Fig. 5), which is the reproductively active adult stage (Fig. 1). These transcripts mainly included the zinc finger domain containing transcripts that are likely acting as transcription regulators. Interestingly, a putative argonaute 2 gene, which is responsible for the defense against viruses and transposons through the RNAi pathway (Zhou and Rana, 2013), was upregulated only in post-aestivation, suggesting a specific need for these functions at this reproductively active stage.

Most of the top DE transcripts related to feeding activity, such as trypsin and other proteinases, were upregulated during pre-aestivation and downregulated during aestivation (Fig. 5). The upregulated transcripts associated with digestive activity differed between pre-aestivation and post-aestivation, which might be reflecting qualitative (different oilseed rape ages) and quantitative (pre-aestivation feeds more than post-aestivation) differences in the feeding patterns of the two stages. Intriguingly, some of these transcripts reached their expression peaks during aestivation, including the cathepsin L-like proteinase, suggesting functions beyond feeding, such as a lysosome-related function.

## 5 DISCUSSION

Aestivation represents an obligatory stage in the life cycle of *P. chrysocephala*, a key pest of winter oilseed rape in Europe (Tixeront et al., 2024; Willis et al., 2020). Despite its importance, the phenomenon of aestivation remains poorly understood, not only in *P. chrysocephala* but also in other insect pests, particularly at the molecular level. This study marks the first comprehensive characterization of the aestivation stage in *P. chrysocephala,* examining both physiological and transcriptional aspects. We identified notable changes in cellular metabolism, mitochondrial activity, reproductive maturity, and body composition in relation to the initiation, maintenance, and termination of aestivation.

The onset of diapause is generally associated with a substantial and sustained decrease in metabolic rate (Lehmann et al., 2020; Melicher et al., 2024; Sgolastra et al., 2010). The fact that aestivation coincides with high temperatures implies that the environmental conditions do not aid the suppression of metabolism and thus might involve mechanisms that have not been described in winter-diapausing insects (Denlinger, 2022; Hahn and Denlinger, 2011). Therefore, prior to conducting the other experiments, we periodically measured the VCO_2_ emission of *P. chrysocephala* adults to rationally select the sampling time points. We observed a significant suppression of VCO_2_ levels beginning in 15-day-old adults, which persisted for 25 days (Fig. 1A). This 25-day period was identified as the aestivation period in our *P. chrysocephala* laboratory population. Notably, this duration was shorter than that reported in a previous study, which documented an aestivation period ranging from 48 ± 17 to 62 ± 14 days (Sáringer, 1984). We attribute this variability in reported aestivation periods to both genetic and methodological differences; notably in the study by Sáringer (1984) the criteria for defining the aestivation period did not rely on metabolic rate and were not precisely defined.

The results from the GO term enrichment analysis align well with the observed metabolic suppression during aestivation, as numerous mitochondria-related terms were enriched in the pre-aestivation vs. aestivation comparison (Fig. 4A). Although VCO_2_ serves as an indirect indicator, primarily reflecting Krebs cycle activity rather than direct measurements of electron transport chain and ATP production, its suppression suggests metabolic alterations. Further investigations showed downregulation of core components of the Krebs cycle, such as NADH dehydrogenase subunits, during aestivation (Fig. 5). Hence, considering both the VCO_2_ and RNA-seq results, we postulate a decrease in the mitochondrial membrane potential and ATP-ADP ratio in *P. chrysocephala* aestivation. These alterations likely facilitate successful aestivation by reducing the demand on internal energy reserves, similar to the diapause in *L. decemlineata*, where mitophagy supports metabolic suppression (Lebenzon et al., 2022). According to the hygric hypothesis, suppression of metabolism indicates reduced respiration rates, which, in other species, has enabled a reduction in water loss rates (Gibbs et al., 2003; Matthews and White, 2012). The enhanced survival under summer conditions might partly be due to the reduction in desiccation risk conferred by the lower respiration rates as well as the upregulation of cytochrome P450s, such as 4C1 (Shen et al., 2021), in *P. chrysocephala*.

During aestivation, *P. chrysocephala* beetles do not feed. However, they resume feeding after aestivation, albeit at a reduced rate compared to their pre-aestivation levels (Ankersmit, 1964; Sivčev et al., 2016). Consistent with these observations, our enrichment analyses indicate a decrease in digestion during aestivation (Fig. 4A, 4B). The downregulation of genes associated with digestion likely represents one strategy to reduce energy expenditure on temporally unnecessary processes, a phenomenon observed, for example, in *Culex pipiens* (Robich and Denlinger, 2005), *G. daurica* (Ma et al., 2019) and *L. decemlineata* (Yocum et al., 2009). Interestingly, many of the genes that were downregulated during aestivation did not return to their pre-aestivation levels upon emergence from aestivation. Additionally, some genes were uniquely upregulated in post-aestivation (Fig. 5), which might be an adaptation to the change of diet from old to young oilseed rapes (Ankersmit, 1964).

The absence of feeding and the corresponding downregulation of digestion genes in aestivating beetles prompts the question: how do these beetles secure the requisite energy for sustaining their basal metabolism during this period? Consistent with the notion that diapausing insects draw upon their internal energy reserves (Denlinger, 2022; Hahn and Denlinger, 2007), we found evidence for metabolic alterations or the mobilization of lipids and carbohydrates during the transition into and out of aestivation at the transcriptional level (Fig. 5B). Moreover, the measured changes in lipid, glycogen, and water-soluble carbohydrate (WSC) levels also indicate that the energy reserves of the adult beetles depend on their diapause stage (Fig. 3B-3D). While glycogen and WSC levels were almost completely depleted during early-mid aestivation, presumably to fuel the basal metabolism, lipid levels were highest during aestivation and were depleted by post-aestivation. This suggests that lipids may be used as a metabolic fuel during aestivation. Similarly, aestivating *Eurygaster maura* males upregulate a lipolytic gene, lipid storage droplet 1, presumably for the same purpose (Toprak et al., 2014). Additionally, one of the cathepsin L-like proteinases emerged as a top-upregulated gene during aestivation (Fig. 5). This proteinase might play a role in inducing matrix metalloproteinases, leading to the degradation of extracellular matrix proteins. This process could potentially trigger fat body cell dissociation and activate lipid metabolism during aestivation. This cathepsin L-like proteinase function was observed, for example, in the pupae of the non-diapausing moth *Helicoverpa armigera* (Jia and Li, 2023). Post-aestivating adults, especially females, may heavily rely on these lipid reserves for oviposition and migration behavior (Sinclair, 2015; Sinclair and Marshall, 2018; Wolda and Denlinger, 1984). This hypothesis is supported by the observation that many lipid metabolism-related genes were uniquely upregulated in post-aestivating individuals (Fig. 5).

Previous studies have indicated that adult *P. chrysocephala* reproduces after completing aestivation (Alford, 1979; Vig, 2003). However, a comprehensive investigation is still lacking. In this study, we observed that females slowly emerged from aestivation after 35 days, as their VCO_2_ levels increased. Yet, females did not have mature ovaries and did not oviposit until they were 45 days old (Fig. 1B, 1D). Likewise, most of the transcripts related to reproduction were upregulated specifically during the post-aestivation phase (Fig. 5).

Numerous transcriptional alterations linked to aestivation cannot be fully explored here due to space limitations. Noteworthy changes include the upregulation of several genes encoding transcription repressors (e.g., zinc finger proteins) and positive transcriptional regulator genes during aestivation compared to the pre-aestivation state. These changes likely contribute to the establishment of the aestivation phenotype through the transcriptional regulation of other relevant genes, a phenomenon observed in other diapausing insects (Bao and Xu, 2011; Guo et al., 2018; Torson et al., 2023; Zhang et al., 2020). Importantly, one specific transcription factor, annotated as SOX-8, showed a unique and substantial upregulation solely during aestivation. This gene potentially serves as a regulator of the genes that facilitate the aestivation phenotype. Future studies should prioritize its characterization to elucidate its role in mediating aestivation-related gene regulation.

The potential functions of aestivation may include synchronization with the life cycle of host plants, avoidance of predators or parasitoids, and resistance to abiotic factors associated with summer, such as high temperature and low humidity (Masaki, 1980; Saulich and Musolin, 2018). *P. chrysocephala* adults are small in size (3-4 mm) (Edde, 2022) and consequently have large surface-area-to-volume ratios, suggesting an increased risk of desiccation during the summer season (Kühsel et al., 2017). Our current study provides evidence supporting the idea that a critical function of aestivation is the mitigation of increased desiccation risk. First, the survival rate in the aestivating *P. chrysocephala* adults was higher than that of pre-aestivating adults. A similar increase in survival under high temperatures was also observed in the diapausing pupa of *Helicoverpa assulta* compared to non-diapausing stages (Liu et al., 2016). Moreover, the observed changes in the transcription of trehalose transporter genes during aestivation (Fig. 4A) may be necessary for maintaining an optimal intracellular concentration of trehalose. This sugar can act as a chemical chaperone against various stress conditions, including extreme temperatures and desiccation (Tang et al., 2008). In addition, it serves as a significant source of energy for sustaining basal metabolism, which produces metabolic water (Kikawada et al., 2007; Zhou et al., 2022).

Several transposon-related transcripts were specifically upregulated during aestivation (Fig. 5). Although similar observations have been made in various diapausing insects, such as *L. decemlineata* (Lebenzon et al., 2021), *C. pipiens* (Robich et al., 2007), and *Drosophila montana* (Kankare et al., 2016) the functional roles of these upregulated transposon-related transcripts during aestivation remain to be elucidated. Indeed, there is an intriguing possibility that transposable elements are involved in the regulation of diapause/aestivation by influencing epigenetic regulatory mechanisms. An alternative explanation could be that diapausing insects are simply unable to inhibit transposon replication due to the shutdown of defense mechanisms, including RNAi pathways, as a means to maintain energy balance. This idea is supported by increased viral RNA copy numbers observed under diapause-inducing conditions compared to non-diapause-destined conditions in *Aedes Albopictus* (Zhang et al., 2022), hinting at a compromised defense against invasive nucleic acids.

Overall, the findings presented in this study, in conjunction with earlier studies (Ankersmit, 1964; Bonnemaison & Jourdheuil, 1954; Sáringer, 1984), provide a comprehensive understanding of the adult stage of *P. chrysocephala.* Initially, emerging adults engage in feeding behavior to accumulate energy reserves such as sugars and lipids. Then, the beetle’s transition into aestivation is presumably facilitated by alterations in hormone titers such as juvenile hormone and specific transcriptional factors. During aestivation, cells downregulate unnecessary processes, including the production of digestive enzymes, and significantly suppress metabolism by modifying mitochondrial and other cellular activities. To sustain their basal metabolism, beetles heavily rely on their sugar reserves such as WSC and glycogen during the early phase of aestivation and then β-oxidize previously accumulated lipid reserves. Upon completing aestivation, beetles further deplete their lipid reserves and resume feeding to reach reproductive maturity and colonize newly sown oilseed rape fields. Future studies should prioritize investigating the importance of each of the highlighted biological processes in the progression, sustenance, and cessation of aestivation. This could be achieved through loss-of-function experiments targeting core genes associated with each respective process.

## CONCLUSION

This study marks the first investigation into the obligatory aestivation process of *Psylliodes chrysocephala* at the mRNA level, complemented by physiological data. The results suggest that newly emerged *P. chrysocephala* adults prepare for aestivation by accumulating energy reserves and subsequently reducing their metabolic rate through alterations in mitochondrial activity and the expression of genes coding for metabolic enzymes. This adaptation allows the insect to compensate for the lack of feeding during aestivation. Conversely, the termination of aestivation is associated with changes in lipid metabolism and transcription factors, enabling the insect to reach reproductive maturity. Notably, our findings have identified several testable hypotheses concerning the key genes and processes underlying the transitioning into and out of aestivation in *P. chrysocephala.* Such insights hold promise for the enhancement of RNAi-based management strategies targeting this pest.

## ACKNOWLEDGEMENTS

The authors thank Daniel Veit (Max Planck Institute for Chemical Ecology, Jena, Germany) for providing the modified LI-820 CO_2_ Gas Analyzer. They also extend their gratitude to Dr. Johan Zicola, Katharina Kubon, Jonas Watterott, and Ruth Pilot for their support. The authors acknowledge the funding provided by the Deutscher Akademischer Austauschdienst (grants to Gözde Güney and Doga Cedden). The graphical abstract was created using BioRender.com, with assistance from Metin Cedden.

## AUTHOR CONTRIBUTIONS

**GG:** Conceptualization, Methodology, Investigation, Data Curation, Formal analysis, Visualization, Writing - Original Draft. **DC:** Methodology, Formal analysis, Software, Data Curation, Visualization, Writing - Review & Editing. **JK:** Methodology. **BU:** Supervision, Writing - Review & Editing. **FB:** Conceptualization, Supervision, Resources, Writing - Review & Editing. **SS:** Conceptualization, Supervision, Resources, Writing - Review & Editing. **MR:** Conceptualization, Resources, Supervision, Writing - Review & Editing.

