## Supplementary information for "Physiological and transcriptional changes associated with obligate aestivation in the cabbage stem flea beetle (*Psylliodes chrysocephala*)"

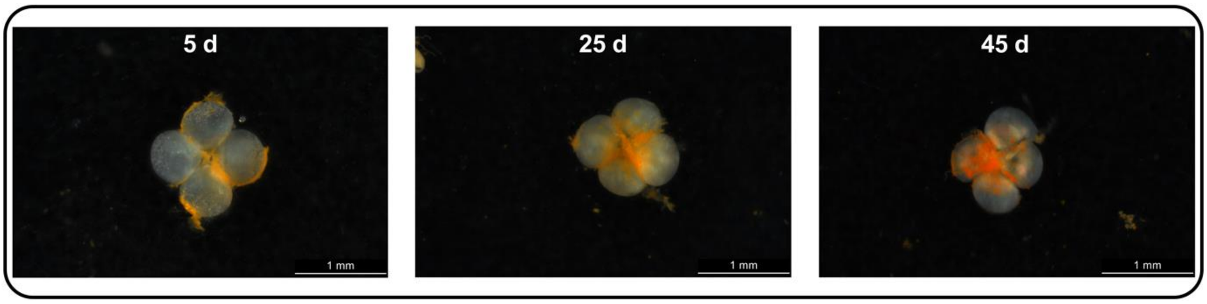


Figure S1. Representative pictures of testes dissected from adult *Psylliodes chrysocephala* males 5, 25, or 45 days post-emergence (n = 5) after the removal of the membranous sac.

Table S1. *P* values resulting from Two-way ANOVA on the body composition investigations.

| | **Stage factor** | **Sex factor** | **Interaction** |
|---|---|---|---|
| Protein content | .078 | .009 | .439 |
| Total lipid content | <.001 | .123 | .094 |
| Glycogen content | <.001 | .759 | .900 |
| WSC | <.001 | .038 | .903 |
| Water % | <.001 | .251 | .089 |

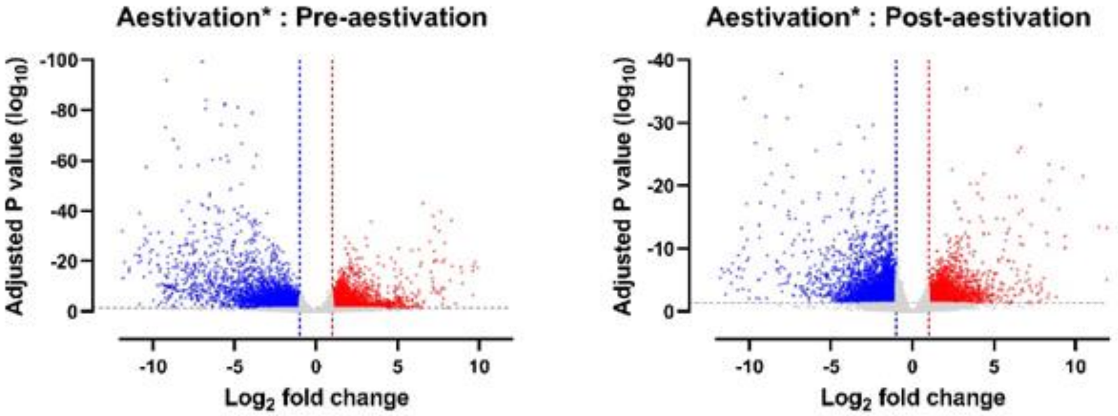


Fig. S2. Volcano plots analyses of differential gene expression analysis

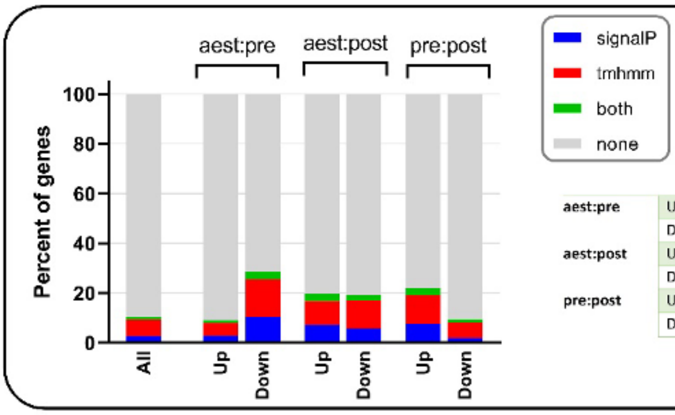


P values

| | | signalP | tmhmm | both |
|---|---|---|---|---|
| aest:pre | Up | .49 | .003 | .21 |
| | Down | <.001 | <.001 | <.001 |
| aest:post | Up | <.001 | <.001 | <.001 |
| | Down | <.001 | <.001 | <.001 |
| pre:post | Up | <.001 | <.001 | <.001 |
| | Down | <.001 | .13 | 0.46 |

Fig. S3. Proportion of signal peptides and transmembrane helicases among the differentially expressed transcripts
